# rgenesconverged : An R Package for the Exploration of Molecular Convergent Evolution

**DOI:** 10.1101/858076

**Authors:** Dina Issakova

## Abstract

Sequence convergence is a type of convergent evolution that results in similarity between orthologous genetic sequences. However, this can be difficult to evaluate statistically given classical paradigms for convergent evolution, which rest on the assumption that convergent phenotypes are achieved by unrelated genetic mechanisms. While sophisticated models exist to model this process and to evaluate the probability of this explanation of genetic similarity across two different species, no tool has currently been implemented in R to carry out these analyses. Here we present **rgenesconverged**, an R package allowing the exploration of sequence convergence hypotheses in a given phylogeny.

## 1. Introduction

Environmental challenges impose on species an unrelenting pressure to adapt, and nowhere is this more evident than in the study of convergent evolution. Contrary to traditional assumptions of divergence, convergent evolution is the evolution of similar phenotypes in response to these challenges by species from independent evolutionary lineages [1]. In one of the most well-known examples, insects, bats and birds have convergently evolved wings with similar characteristics, giving them the ability to fly to escape predators and pursue prey [2]. In this way, environmental conditions select for similar evolutionary solutions to a given problem. Historically, research on convergent evolution has focused on macro-phenotypes: traits clearly observable without a descent to the genome level. In these cases, we assume that convergent evolution has occurred if we observe similar traits without a common genetic basis; it is not the same genetic pathway, for example, that is responsible for flight in birds and mammals [3]. The situation becomes more complicated, however, when we consider ‘micro-phenotypes’, or phenotypes occuring at the protein level. It may not be biologically possible to have sufficiently distinct genetic mechanisms to produce very similar proteins to perform very similar functions in a cell. In such a case, two similar proteins or protein motifs can arise independently in separate lineages and have a similar genetic code without having come from a common ancestor, rendering the traditional test for sequence dissimilarity invalid. This is referred to as sequence convergence, contrasting with functional, mechanistic, and structural convergence [4]. How do we determine whether convergent evolution has occurred in such a case?

This question has been addressed with mathematical models, beginning with the simplest principle: a definition of convergence for micro-phenotypes that requires a genetic similarity between species and a dissimilarity from a common ancestor [5]. When the ancestor is unknown, this method relies on probabilistic reconstruction, commonly using parsimony or maximum-likelihood models of phylogenetic history [6], [7]. Other methods continued to build on the statistical rigour of the earlier methods [8]. Modern methods for the assessment of sequence convergence quantify a nuanced convergence score based on the similarity between two sequences, based on similarity matrices such as BLOSUM62. Then, the task becomes to convince ourselves that the similarity between two sites is not due to common descent from a molecule with the same function or feature [9]. This comes principally from the comparison to orthologs in related species. It is crucial to consider the chance of the observed convergence event occuring by chance; after all, there are only 21 amino acids, and the genetic code at any locus is transformed over time by mutation and genetic drift. The statistical methods described above were devised with this rigour in mind, allowing us to differentiate convergence from random sequence similarity [5][10]. In experimental work it is also crucial to consider how related the sequence of interest is to its function, as well as the possibility of horizontal gene transfer or recombination.

While this remains an active area of research, standard statistical modesls for sequence convergence have been established in the field and have since been used in a variety of publications [11]–[14]. At the same time, bioinformatics tools have become more and more important in evolutionary research [15]–[17]. Particularly, there has been an emphasis on data analysis in R [18], [19]. While multiple R packages exist to parse, reconstruct and analyse genes in a phylogenetic context, no package has been developed dedicated to convergent evolution that implement the common models used in the field. With this in mind and building on top of the existing tools [20]–[22], we have developed rgenesconverged, an R package that implements these models for ease of use by researchers interested in sequence convergence, and extends existing models to allow for exploratory computational work. We provide extensive documentation, a graphical user interface for users unfamiliar with R, and an example workflow walk-through.

## 2. Setup and Workflow Overview

The inputs required to rgenesconverged are an R phydat object containing sequence information for the selected species and an R tree object. Multiple tools exist to convert trees in standard forms such as Newick into this format, provided for example by the package ape [22]. The user selects a species of interest for comparison to which the other species in the tree are compared. Sequences are aligned by MUSCLE to the reference sequence; the position at which convergence is to be evaluated corresponds to that position in the reference sequence, and corresponding positions in the alignment.

Two possible ‘paths’ exist in this package for phylogenetic analysis. The first is a direct implementation of Zhang and Kumar’s model [5], which is a well-established and commonly used model for sequence convergence; the second is a natural extension allowing for a more nuanced and exploratory analysis, but less statistical rigor. An R plot is also available, which finds species that are convergent at the particular position specified to the reference species, and highlights them in a phylogram.

While designed for use in an R environment, an RShiny UI is available with the package for users unfamiliar with R programming.

All code for rgenesconverged is available at **https://rdrr.io/github/dinaIssakova/rgenesconverged**.

### Note on preprint

As this project is still in active development, updates are continually made to the github repository. It is available under the MIT licence and the software is provided ‘as is’ without warranty.

## 3. Zhang and Kumar: Conditions for Convergence

As described by Zhang and Kumar [5], an amino acid position is considered convergent across two separate lineages if the two are identical and different from their common ancestor (while the original definition was that it must differ at some point across a lineage, in practice this often does not match up with phylogenies analysed in R, thus leading to this slightly generalised definition).

Of course, this leads to an obvious caveat - suppose we have two amino acids, *a* and *x*, which are very similar to one another and frequently intermutate. Two present-day species have amino acid x at some position in the same locus, while their common ancestor had a at the same position. Is this evidence of convergent evolution? By this definition, yes; however, there is a high likelihood, especially if the two genes’ coalescence point is far in the past, that this has simply occured by chance. For this reason rgenesconverged reports *p* values for all calculations, and reports instances of convergence as potential events.

## 4. Probability of Convergence

**Figure 1:**
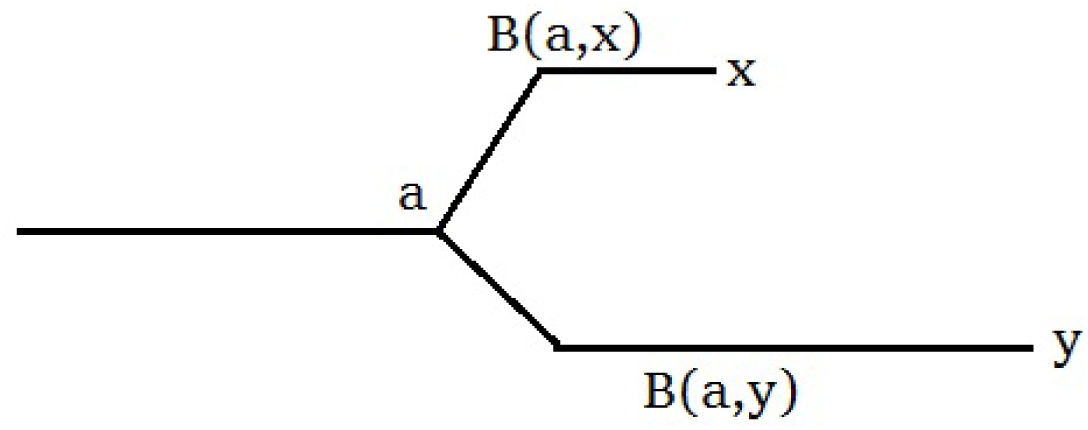
A general example provided for the statistical discussion.

As described by Zhang and Kumar, we can describe the probability of our observation being a chance occurence in the following way.

Following our earlier example, suppose we call our modern-day amino acid states *x* and *y*, sharing an ancestral state *a*, with corresponding branch lengths of *B*(*a, x*) and *B*(*a, y*) respectively.

The probability of a site mutating, say from *a* to *x* over branch length *B*(*a, x*), *P*_*ax*_(*B*(*a, x*)) can be computed by a typical similarity matrix such as PAM or BLOSUM [23]–[25] (PAM is used here as it was referenced in the original publication [5]). The probability of a given site configuration then becomes the probability of the occurrence of *a* (defaulted as 1/20, but can be adjusted by the user), multiplied by the likelihood of *a* mutating to *x* and *y* over their respective branch lengths. Since we are only interested in the probability of configurations satisfying the requirement that *x* = *y* and *x, y* ≠ *a* we then find the probability of any configuration satisfying the hypothesis of convergent evolution under the constraint of the given tree, defined as *f*, as

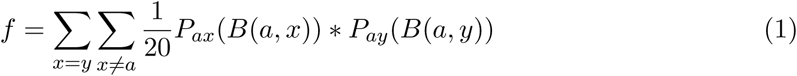

This eventually [5] leads to the following conclusion: that if a sequence is *m* amino acids long and assuming a uniform substitution rate, the probability of *n* convergent sites (*ϕ*) using the definition above is

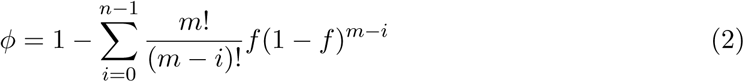

Knowing this metric, we can easily distinguish whether or not our convergence hits are meaningful. This is applicable when n is greater than or equal to 1; if n = 0, then *ϕ* = 0. In some cases, this can be approximated with a Poisson distribution [5], but this is not implemented in rgenesconverged as of writing time.

## 5. Extending the Model - General convergence score

**Figure 2:**
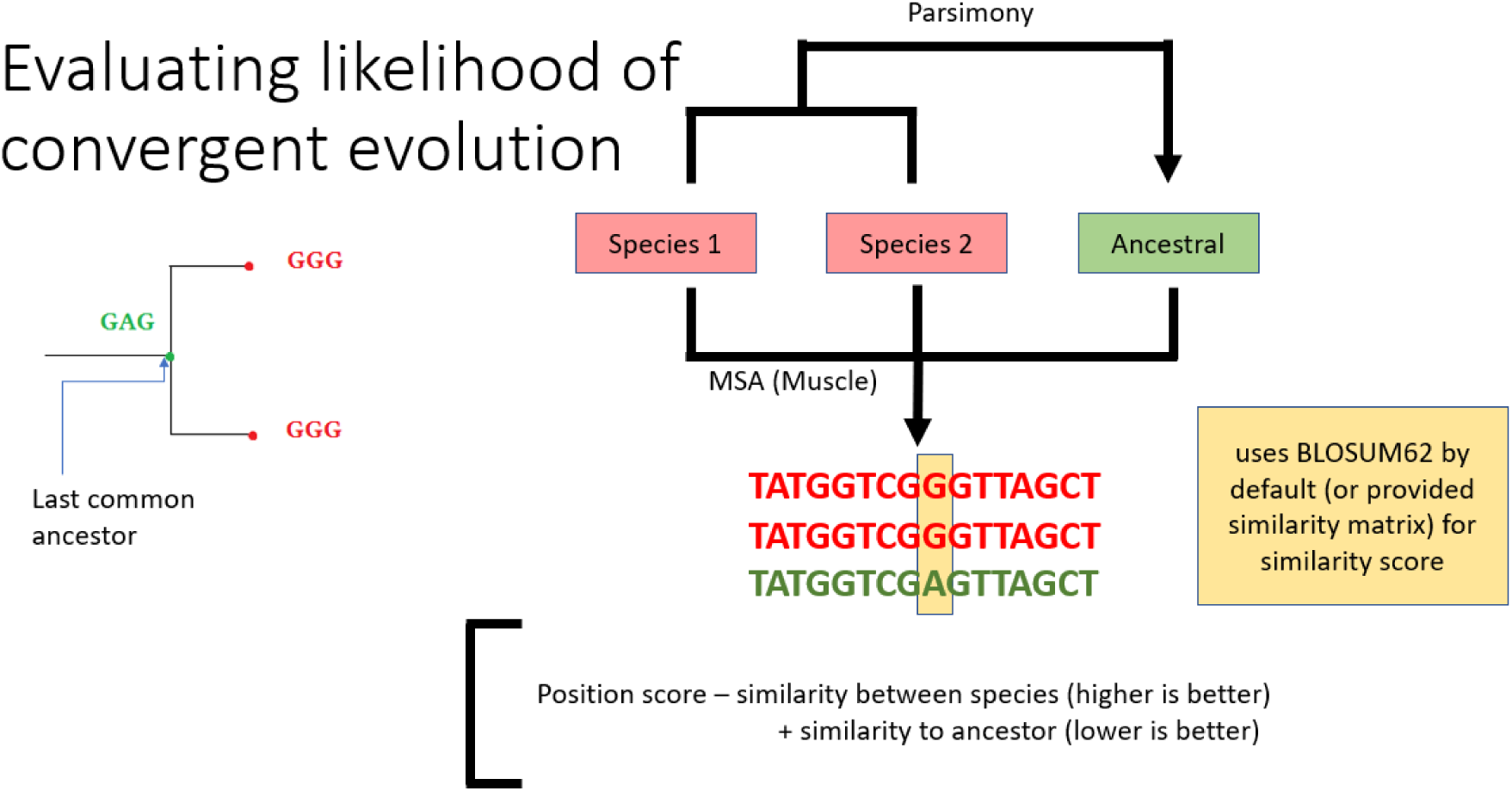
The general workflow of scoring a convergent site.

To provide a less deterministic alternative more suitable to interactive exploration, we have implemented an alternative scoring method for convergence into rgenesconverged. All positions are relative to the species selected by the user; the positions of the other species are the amino acid present at the aligned position. The total convergence score at a given positon consists of two parameters. The first is the similarity score given by BLOSUM62 between the two amino acids at the respective position in the two sequences: it is higher when the sequences are more similar. The other is a measure of the dissimilarity of the species of interest from the ancestral state, which is obtained by taking the negative BLOSUM62 similarity score: the more different the amino acid is from its ancestral state, the more confidently can we claim conversion.

Thus, the total score is equal to the similarity between the reference species and the ancestor subtracted from the similarity between the two modern-day lineages.

## 6. Evaluating probability of null hypothesis

Similarly to the model devised by Zhang and Kumar, we need to calculate that our observations are due to chance. In this case, we calculate the probability of a configuration having a convergence score greater than the specified threshold, over all possible values of *x, y, a*.

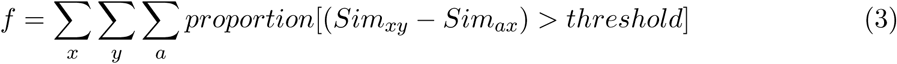

In the same way as above, we substitute the value of *f* into equation 2 to get the probability of *n* or more convergent sites (in this case, sites that score above the threshold) in a sequence.

## 7. Conclusion

While rgenesconverged is a package still in active development, this preprint is provided with the hope that it can provide useful functionality to researchers studying convergent evolution. This package provides a much-needed implementation of a classic convergence model to be used in R, as well as a UI for users without programming experience. It also extends the model, adding a scoring measure to allow for more nuanced exploratory analysis. Future and current work on this package involves adding alignment visualisation into the phylogenetic tree representation, as well as extending the UI to fully cover all aspects of the package available to R users. We wish also to extend our statistical model to allow for cases where no ancestral value exists, for example for convergence as it applies to insertions or deletions.

Including both in-text citations and packages whose functions were used in building rgenesconverged.

